# Non-invasive Chromatin Deformation and Measurement of Differential Mechanical Properties in the Nucleus

**DOI:** 10.1101/2021.12.15.472786

**Authors:** B. Seelbinder, M. Jain, E. Erben, S. Klykov, I. D. Stoev, M. Kreysing

**Affiliations:** Max Planck Institute of Molecular Cell Biology and Genetics, Dresden (Germany); Centre for Systems Biology, Dresden (Germany); Cluster of Excellence Physics of Life, TU Dresden, Dresden (Germany)

**Keywords:** Nuclear Mechanics, Chromatin Organization, Thermophoresis

## Abstract

The nucleus is highly organized to facilitate coordinated gene transcription. Measuring the rheological properties of the nucleus and its sub-compartments will be crucial to understand the principles underlying nuclear organization. Here, we show that strongly localized temperature gradients (approaching 1°C /μm) can lead to substantial intra-nuclear chromatin displacements (>1 μm), while nuclear area and lamina shape remain unaffected. Using particle image velocimetry (PIV), intra-nuclear displacement fields can be calculated and converted into spatio-temporally resolved maps of various strain components. Using this approach, we show that chromatin displacements are highly reversible, indicating that elastic contributions are dominant in maintaining nuclear organization on the time scale of seconds. In genetically inverted nuclei, centrally compacted heterochromatin displays high resistance to deformation, giving a rigid, solid-like appearance. Correlating spatially resolved strain maps with fluorescent reporters in conventional interphase nuclei reveals that various nuclear compartments possess distinct mechanical identities. Surprisingly, both densely and loosely packed chromatin showed high resistance to deformation, compared to medium dense chromatin. Equally, nucleoli display particularly high rigidity and strong local anchoring to heterochromatin. Our results establish how localized temperature gradients can be used to drive nuclear compartments out of mechanical equilibrium to obtain spatial maps of their material responses.

**Main Findings:**

- Novel non-invasive active micro-rheology method to probe spatial intranuclear material responses, unhindered by the nuclear lamina, using strongly localized temperature gradients
- Chromatin shows both elastic and viscous properties at the mesoscale with a retardation time of τ ∼ 1s
- Compacted heterochromatin in a model of nuclear inversion shows high resistance to deformation, suggesting dominantly solid-like behavior
- The nucleus displays spatially distinct material properties for different compartments
- The nucleolus shows high resistance to deformation on the time scale of seconds
- Immobile nucleoli appear solidly anchored to and retain the deformation of surrounding chromatin

## INTRODUCTION

It is widely believed that the spatial organization of the nucleus is supported by and makes functional use of distinct material properties to support homeostatic function (Cremer et al., 2020; Falk et al., 2019; Mirny and Dekker, 2021), and that may be temporally adapted to facilitate cell cycle dynamics and differentiation (Mittasch et al., 2020; Strom et al., 2021; Sun et al., 2018). Starting at the nanometer scale, molecular interactions are thought to give rise to the spatial organization of nuclear constituents that become visible at the micrometer scale. For example, the nucleus features membraneless organelles, such as the nucleolus, Cajal bodies, nuclear speckles, PML bodies and others, which are comprised of RNA and proteins and are considered to be formed by liquid-liquid phase separation (LLPS) (Zidovska, 2020a). (Feric et al., 2016). Most of the nucleus is occupied by chromatin, however, which is hierarchically organized into different compartments: i) a few nucleosomes (5-20) loosely assembled into clutches, which further assemble into chromatin nanodomains, ii) nanodomains are further grouped into local continuous gene clusters called topologically associated domains (TADs), and iii) TADs from different loci group together to form two main compartments: active A-compartments and inactive B-compartments (Jerkovic’ and Cavalli, 2021; Mirny and Dekker, 2021; Misteli, 2020). Using data from chromatin conformation capturing assays (Hi-C), computer simulations suggest that strong interactions between B constituents together with weak interactions of A constituents drive the separation of compartments (Falk et al., 2019). Furthermore, condensed chromatin is thought to solidify with increasing strength of molecular interactions, yielding elastic rather than viscous responses (Hansen et al., 2021). Hence, differences in interactions within compartments should also be directly measurable as a reflection of different material properties. Experimental characterization of the material properties of nuclear compartments will be crucial to understand spatial nuclear organization and its role in nuclear information processing. Despite great progress, an integrated physical picture of the nucleus is still missing.

A recent point of discussion has been whether chromatin behaves like a liquid or a solid (Strickfaden et al., 2020; Zidovska, 2020b). For chromatin it has been suggested that its material properties are predominantly liquid in line with the view that heterochromatin domains form through LLPS, facilitated through the binding to scaffold proteins, similar to nuclear organelles (Gibson et al., 2019; Larson et al., 2017; Strom et al., 2017). However, FRAP experiments in interphase nuclei that harbored fluorescently-labeled chromatin observed that bleached hetero- or euchromatin regions did not recover their intensity after bleaching at the minute to hour scale (Strickfaden et al., 2020). Contrary to the earlier view, this indicates that chromatin cannot move freely and therefore behaves more like a solid. To reconcile the observed discrepancies, it has been suggested that chromatin, like other polymers, shows a more complex behavior that can be viscous, elastic or viscoelastic depending on the time and length scales that are being probed and the energy-driven enzymatic activity in the environment (Zidovska, 2020b; Zidovska et al., 2013). For example, the liquid-like behavior of chromatin observed at the nanoscale (Nozaki et al., 2017) likely emerges from enzymatically driven processes such as transcription and loop extrusion (Fudenberg et al., 2016; Golfier et al., 2020). A high molecular mobility on the nanoscale might still be locally constrained and prevent large scale rearrangements of chromatin. Hence, observing liquid-like behavior at the nano scale does not have to be in contradiction with solid-like behavior at the micron scale.

One functional adaptation of nuclear architecture is the central compaction of heterochromatin in rod cells, termed nuclear inversion. (Solovei et al., 2009). Nuclear inversion is triggered by LBR down regulation (Solovei et al., 2013) and describes the successive fusion of chromocenters during terminal stages of retinal development, which leads to improved contrast sensitivity under low light conditions (Subramanian et al., 2021). Motivated by these findings, it has been suggested that heterochromatin cohesion drives nuclear inversion, as well as the separation of hetero- and euchromatin in conventional interphase nuclei (Falk et al., 2019). Yet, it remains an open question which material properties compacted heterochromatin adapts, how more heterochromatin is mechanically integrated into the nucleoplasm and how chromatin interfaces with other nuclear compartments. To better understand chromatin organization across scales, new experimental approaches are needed to measure material properties in living cells, ideally spatially resolved and dynamically.

A wide range of different complementary techniques have been proposed to infer compartment interactions or nuclear material properties. At the nanoscale, Hi-C based methods have been used to map the spatial interactions between different chromatin compartments (Belaghzal et al., 2021; Falk et al., 2019; Lieberman-Aiden et al., 2009) and reconstitute the genomic 3D organization in silico (Stevens et al., 2017). While Hi-C reports on genome interactions and allows to infer the spatial organization to a very good degree, Hi-C methods themselves do not capture dynamic processes or facilitate making predictions about the material properties, without being complemented by other techniques.

To acquire dynamic data of chromatin motion at the mesoscale (0.01-1 μm), passive microrheology approaches can be used to infer material properties from the spatio-temporal dynamics of discernable features using video microscopy in conjunction with tracking algorithms (Armiger et al., 2018; Herráez-Aguilar et al., 2020; Nozaki et al., 2017; Zidovska et al., 2013). These methods are successful in gaining insights into apparent material properties non-invasively; however, their use is limited to short time scales during which thermal fluctuation dominates over active processes. At larger time scales, motion appears to be largely driven by active ATP-dependent processes and material properties cannot be quantified anymore by assuming thermal motion as the driver. (Guo et al., 2014; Zidovska et al., 2013). For example, chromatin shows ATP-dependent coherent motion on time scales above one second (Zidovska et al., 2013).

To overcome the limitations of passive micro-rheology in energy-consuming materials, active micro-rheology approaches have been developed that use an external stimulus to drive materials out of their mechanical equilibrium. Methods to test nuclear material properties include AFM (Liu et al., 2014), micropipette aspiration (Davidson et al., 2019), magnetic beads (Guilluy et al., 2014; Keizer et al., 2021) and membrane stretch devices (Schürmann et al., 2016; Seelbinder et al., 2020). A difficulty, however, is that the nuclear lamina, an outer stiff shell that surrounds and protects the nucleus, is believed to be at least 10x stiffer than chromatin and would therefore mask the internal material properties when probed from the outside (Harada et al., 2014; Isermann and Lammerding, 2013). These methods provide a good understanding of the mechanics of the nucleus as an integrated whole, but lack spatial resolution. Recently, the injection of magnetic beads into live nuclei allowed for the estimation of local material properties inside the nucleus, but spatial control of the bead location is still limited (Keizer et al., 2021).

Spatially resolved maps of material properties can be obtained by Brillouin Microscopy (Brillouin, 1922; Prevedel et al., 2019; Scarcelli and Yun, 2008), by quantifying Raman shifts of scattered photons. Brillouin Microscopy (BM) has been used to map mechanical properties in zebrafish (Sánchez-Iranzo et al., 2020) and living cell nuclei (Zhang et al., 2020, 2017). While it is a powerful method, BM measures the mechanics at very short time scales (sub-nanosecond) due to its reliance on acoustic waves in the GHz range. Highly attractive would therefore be a micro-rheology method that permits probe-free, spatially resolved mapping of material properties on physiologically relevant time scales, while leveraging the conceptual advantages of active perturbations.

Here, we report on a novel approach that utilizes highly localized temperature gradients to displace and strain chromatin inside the nucleus. Since displacements can reach hundreds of nanometers to few microns, local displacements can be quantified through tracking algorithms. As temperature stimuli pervade the stiff nuclear lamina, chromatin motion occurs without disturbing nuclear size or shape. Using this method, we observed that interphase chromatin displays highly reversible, viscoelastic behavior in response to deformation with a characteristic time of τ∼1 s. We find that material properties are spatially distinct for different compartments.

The nucleolus, in particular, shows high mechanical resilience to deformation, but significant adhesion to surrounding chromatin.

Together, our results showcase the utility of this new approach to actively and non-invasively probe nuclear material properties in a spatially and temporally resolved manner, enabling to assign material responses to biochemical identity of compartments and gain new insights into the mechanics of compartment interfaces.

## RESULTS

### Engineered temperature gradients facilitate controlled chromatin deformation in living cells

To generate a highly localized temperature gradient, a heating IR-laser beam (λ=1455nm) was focused and rapidly scanned (1 kHz) along a line (**Fig. 1a**). Temperature profiles for various laser intensities (low, medium, high) were measured using the temperature-sensitive dye Rhodamine B at a chamber temperature of 36°C (**Fig. 1b-c, Suppl. Fig. 1a**). The average temperature increase inside the nucleus could further be confirmed via temperature sensitive mCherry-H2b, which corresponded well to Rhodamine measurements averaged along a typical nuclear length of 20 μm (**Fig. 1d-e, Suppl. Fig. 1b**). Unless otherwise indicated, experiments were run at medium laser intensity resulting in an average temperature increase of 2°C inside the nucleus.

**Fig. 1.**
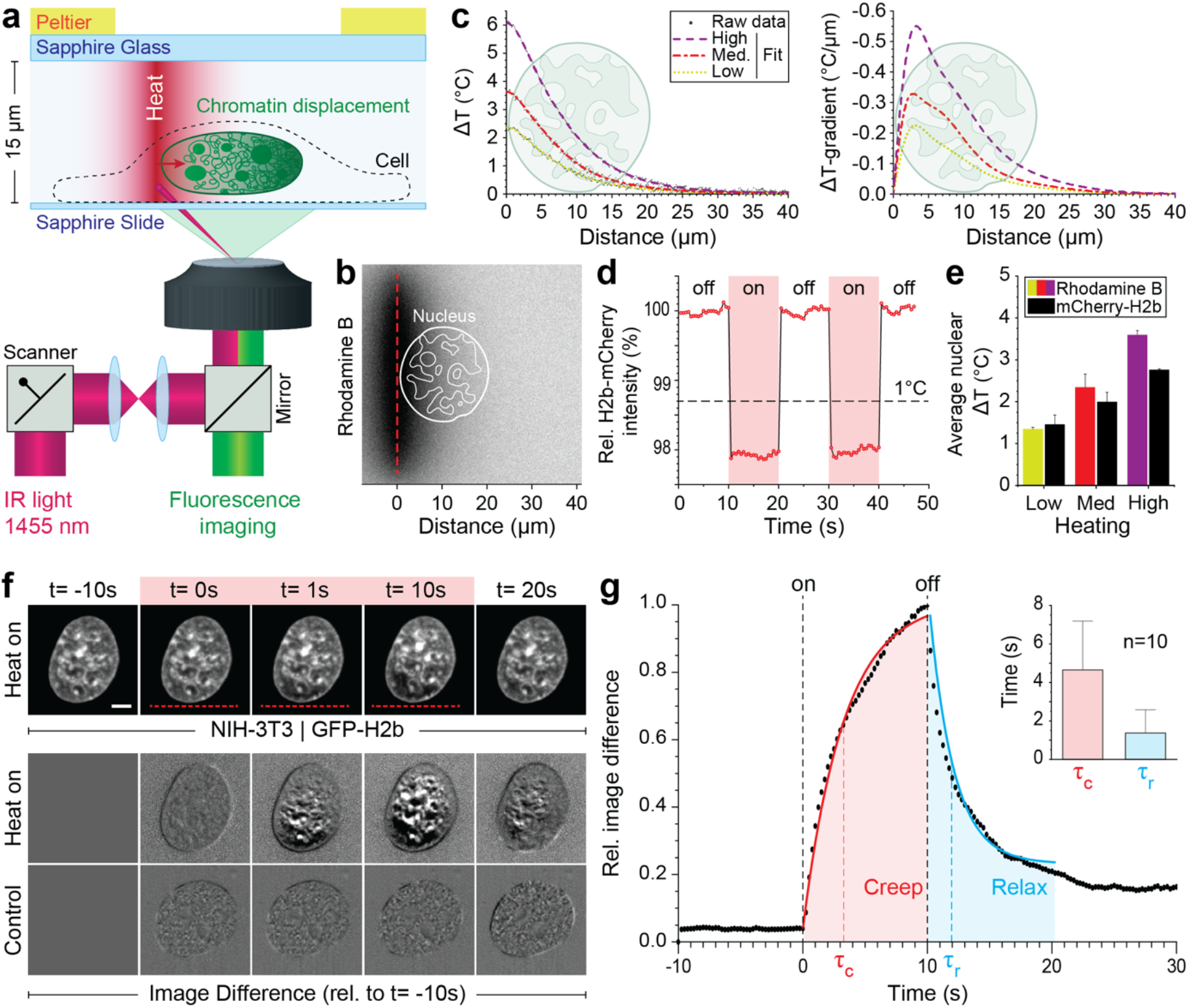
Engineered temperature gradients facilitate controlled chromatin deformation in living cells. **a)** Overview of the microscopy setup. An IR laser is scanned along a line to generate a temperature gradient perpendicular to the scanning line. Cells are cultured in temperature chambers that maintain a reservoir temperature of 36°C via Peltier elements. Sapphire, rather than glass, was used due to its high heat transfer coefficient. Shown is the side view of chamber. **b)** The temperature-sensitive dye Rhodamine B can be used to visualize temperature gradients during laser scanning. The red dotted line indicates the laser scan path and outlines of an average size nucleus are superimposed in white. **c)** Temperature profile and gradient were quantified perpendicular to the laser scan path via Rhodamine B for three different laser intensities (low, medium and high). Raw data of temperature measurements were fitted and differentiated to achieve noise-robust estimates of the temperature gradients. See Suppl. Fig. 1a for dye calibration. **d)** Confirmation of instantaneous temperature changes inside the nucleus, at low laser intensity, via relative quantum efficiency measurements of mCherry-H2b, which reduces ∼1.3% for each 1°C heating. See Suppl. Fig. 1b for calibration. **e)** Comparison of thermometry results measured inside the nucleus via mCherry-H2b and inside the chamber via Rhodamine B. Chamber temperature were averaged along 20 μm, reflecting the average size of a nucleus, to compare both measurements directly; n=5, error=STD. **f)** Top: Response of NIH-3T3 nuclei to an applied temperature gradient at medium laser intensity. Bottom: Image difference analysis of the same data and controls, indicating mesoscopic, partially reversible chromatin displacements due to temperature stimulation. See also Suppl. Videos 1 and 2.m**g)** Temporal analysis of image differences show that chromatin re-arrangements have characteristic times (τ_c_ - creep or retardation time, τ_r_ - relaxation time) are on the time scale of seconds; n=10, error=STD.

When placing the heating stimulus in close proximity to the nucleus, we observed chromatin motion down the temperature gradient as visualized by GFP-H2b in NIH-3T3 cells (**Fig. 1f, Suppl. Videos 1-2**). To better visualize time-dependent chromatin displacements, we generated image difference stacks by subtracting the first image from the image stack (**Fig. 1f**). Analyzing the average image difference over time, we demonstrated that chromatin movement is instantaneous and largely reversible (**Fig. 1g**). Further, image difference dynamics followed an exponential trend that could be fitted to a simple viscoelastic material model (Kelvin-Voigt), with characteristic times τ on the second scale. In the absence of temperature stimulation, we only observed small changes in the image difference stack that likely reflect the spontaneous coherent motion of chromatin as reported before (Zidovska et al., 2013) and might account for some of the residual differences after perturbations.

### Strongly localized temperature-gradients drive intra-nuclear chromatin displacement without affecting nuclear shape

To better characterize the observed chromatin displacement away from an applied temperature gradient, we tracked large intra-nuclear features during the perturbation. Large organelles that exclude chromatin, such as the nucleolus, show up as dark pockets (features) in GFP-H2b images and are well suited to track intranuclear movement (**Fig. 2a**). Kymograph analysis of chromatin-void features confirm instantaneous directed motion with amplitudes of hundreds of nanometers to a few microns, with reversible asymptotic exponential dynamics. (**Fig. 2b-c**). Furthermore, centroid tracking of prominent features reveal that displacements are larger closer to the temperature stimulus.

**Fig. 2:**
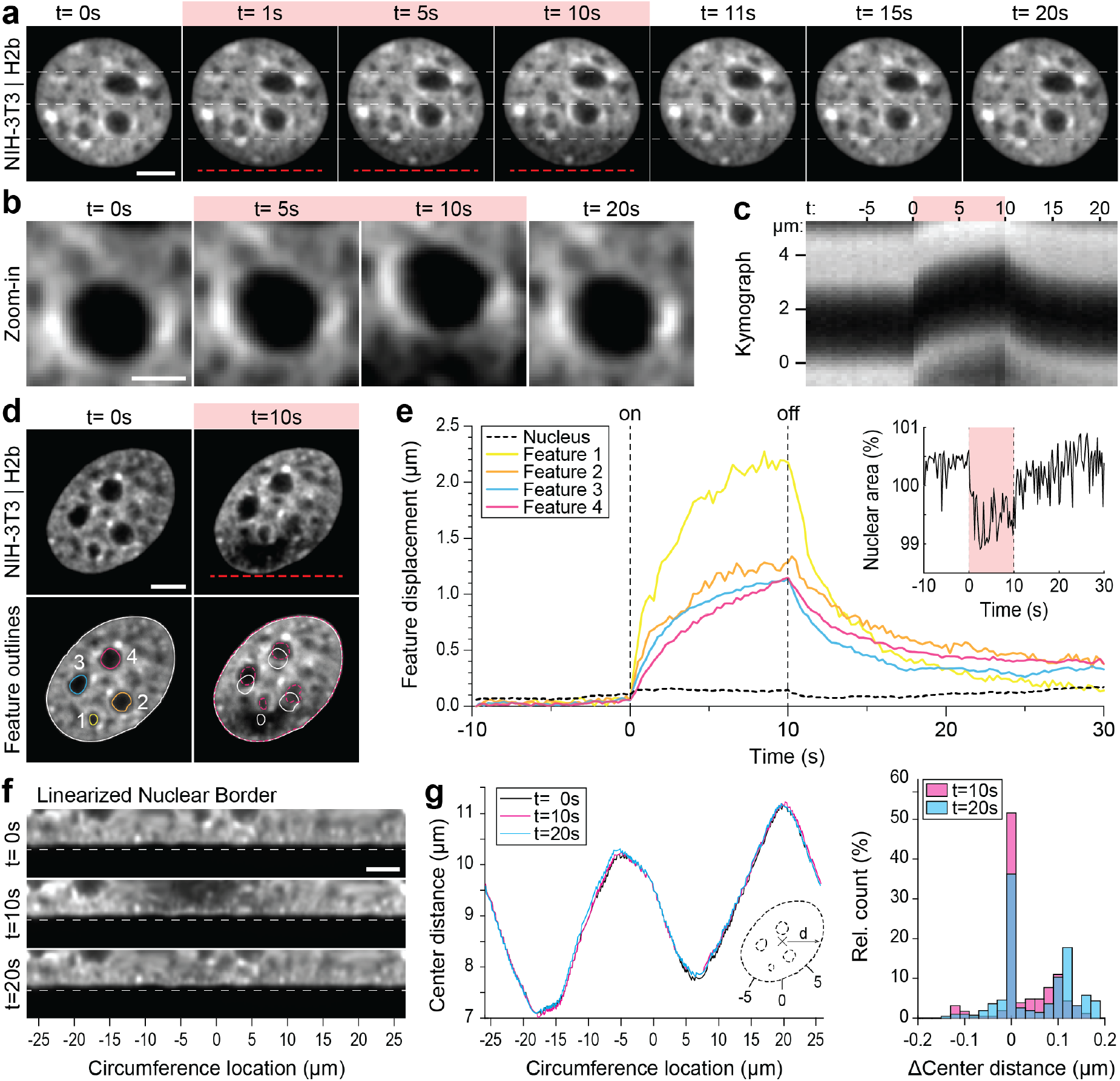
Quantification of intra-nuclear chromatins displacement and absence of changes in nuclear shape during temperature stimulation. **a)** Chromatin displacement over time during 10s temperature stimulation experiments in NIH-3T3 nuclei expressing H2b-GFP; scale=5 μm. **b-c)** Detailed view of one chromatin void feature. Kymograph analysis quantifying time and length scale shows largely reversible motion on the order of microns; scale=5 μm. **d-e)** Segmentation and tracking of intranuclear features show their gradual, heterogenous displacements of up to 2 μm, while nuclear area remains largely constant. See also Suppl. Video 3; scale=5 μm. **f)** Close-up view of the linearized nuclear border of the nucleus shown in d) further indicates that the border remains static during and after temperature stimulation; scale=5 μm. **g)** Quantification of the distance of the nuclear border, with respect to the nuclear center at t=0s, of the nucleus shown in d) reveals that the distortions of the nuclear shape are less than 200 nm during temperature stimulation.

Displacements appear to be restricted to the inside of the nucleus, as the centroid of the nucleus and nuclear area only marginally changed (∼1%) during temperature stimulation (**Fig. 2d-e, Suppl. Video 3**). Specifically, quantifying the change in nuclear geometry by measuring the distance of the nuclear border to the nuclear center before (t=0 s), during (t=10 s) and after (t=20 s) temperature stimulation, we further validated that the deviations in nuclear shape were on the order of 100 nm (**Fig. 2f-g**), similar to displacements of the nuclear centroid. The constant nuclear dimensions likely reflect the dominating stiffness of the nuclear laminar. Additionally, the constant volume of the nucleus distinguishes our findings from the nuclear swelling that was observed during bulk heating of isolated nuclei (Chan et al., 2017).

To summarize our methodological advancement to this point, we found that strongly localized temperature gradients with a magnitude below 2°C cause micron-scale displacements of chromatin and chromatin void organelles inside the nucleus without changing the nuclear area or disturbing the nuclear border.

### Centrally compacted hetero-chromatin behaves as an elastically suspended solid in a genetically induced nuclear inversion model

Tethering of chromatin to the nuclear border (perinuclear chromatin) is an important mechanism that shapes global chromatin structure (Guelen et al., 2008). For example, detachment of chromatin from the nuclear envelope by downregulation of lamin A/C and lamin B receptor leads to the inversion of the conventional chromatin architecture in murine photoreceptor cells with nuclei displaying a condensed heterochromatin cluster in the center (Solovei et al., 2013, 2009). This change in nuclear organization has functional implications even beyond gene expression control, as it serves to improve nocturnal vision in mice (Subramanian et al., 2021, 2019). Due to its characteristic organization, inverted nuclei have become a prominent model to study heterochromatin formation and global genome organization (Solovei et al., 2009).

By overexpressing Casz1 (a zinc finger transcription factor) in NIH-3T3 cells, as shown before (Mattar et al., 2018), we were able to induce chromatin inversion that, in some instances, leads to the formation of a single large central heterochromatin cluster (CHC), reminiscent of the organization of photoreceptor nuclei in mice (**Fig. 3a**). Inverted nuclei present an interesting model for studying chromatin organization, specifically with respect to the material properties associated with heterochromatin formation. We observed a highly reversible displacement of the CHC upon temperature stimulation in videos and kymographs (**Fig 3b, Suppl. Video 4**), which was further verified by tracking of the CHC centroid (**Fig. 3c**). At the same time, area and shape of the CHC remained constant during stimulation, indicating strong resistance to deformation and hence dominantly solid-like behavior (**Fig. 3d**).

**Fig. 3:**
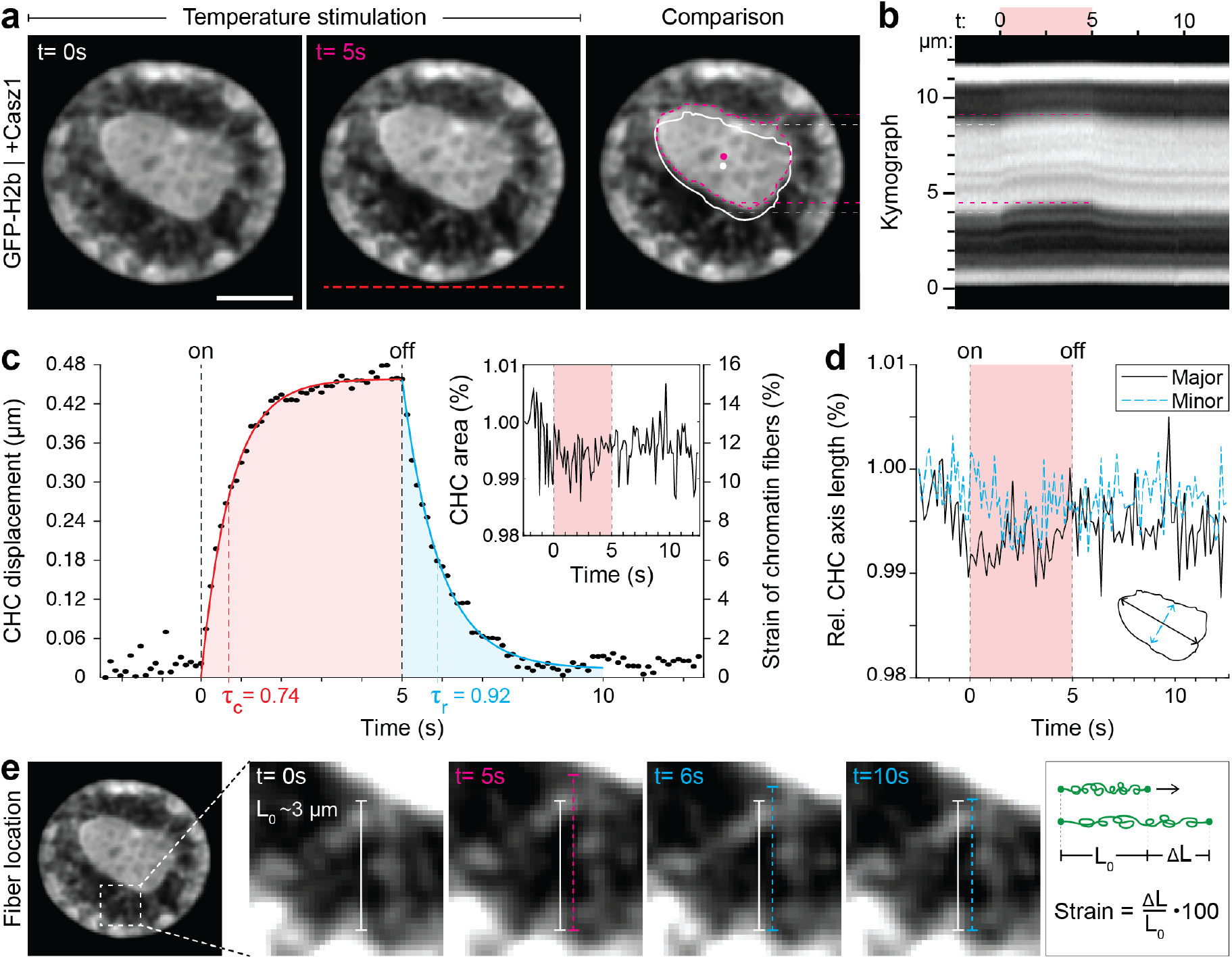
Quantification of movement and shape changes of centrally compacted chromatin during temperature stimulation in a genetically induced nuclear inversion model. **a)** NIH-3T3 cells expressing H2b-GFP displayed an inverted chromatin organization after transfection with Casz1. The resulting central heterochromatin cluster (CHC) shows a significant displacement during temperature stimulation. See also Suppl. Video 4; scale=5 μm. **b)** Kymograph analysis quantifying time and length scale shows reversible submicron scale motion of the CHC. **c-d)** Detailed analysis of the CHC centroid verifies that its motion is characterized by a fast and reversible displacement during temperature stimulation. However, the CHC appears resistant to deformation as it shows little change in area and major and minor axis length. **e)** Close-up view of H2b-positive chromatin fibers straining during temperature stimulation. The cartoon on the right depicts the concept of strain. Based on their initial length L_0_ ∼3 μm, the estimated fiber strain is indicated in c) on the right axis.

The dynamic displacement again fitted well to a simple viscoelastic Kelvin-Voigt model with characteristic times of τ = 0.74s during the creep and τ = 0.92s during the relaxation phase (**Fig. 3c**). Since the CHC moved in its entirety and showed little deformation, this dynamic likely reflects the properties of chromatin fibers that span radially from the CHC to the nuclear border (**Fig. 3e**). By measuring the initial length of the fibers at rest (L_0_∼3 μm) and assuming that the CHC centroid displacement was similar to the extension of fibers (ΔL), the strain of chromatin fibers was estimated to be up to 15% (Fig. 3c, right axis). In contrast, the bulk strain of the CHC was below 1%, indicating that the nucleus features heterogeneous compartment-specific material properties.

### Spatially resolved strain maps reveal distinct mechanical properties of nuclear sub-compartments

Analyzing the chromatin displacement in inverted nuclei provided further evidence that the nucleus features spatially distinct heterogenous nuclear material properties. Therefore, we asked if we could map intranuclear strain more generally in conventional interphase nuclei in order to correlate local material identities with biological function. To this end, we used PIV (Sveen, 2004; Zidovska et al., 2013) to generate spatial displacement maps during temperature stimulation using the frame at t=0s as reference (**Fig. 4a, Suppl. Fig. 2**). To further estimate local intra-nuclear deformation, strain maps were calculated from displacement maps. Shown here are local volumetric changes (hydrostatic strain) and orthogonal displacements (shear strain). Absolute nuclear strain magnitudes, integrated over the whole nucleus, displayed asymptotic exponential dynamics similar to tracked features before (**Fig. 4b**). Averaging non-absolute (total) hydrostatic strains over the whole nucleus, where positive values (extension) and negative values (compression) can cancel each other, results in a line close to 0%, which is congruent with our observation that there is little change in overall nuclear area during temperature stimulations.

**Fig 4:**
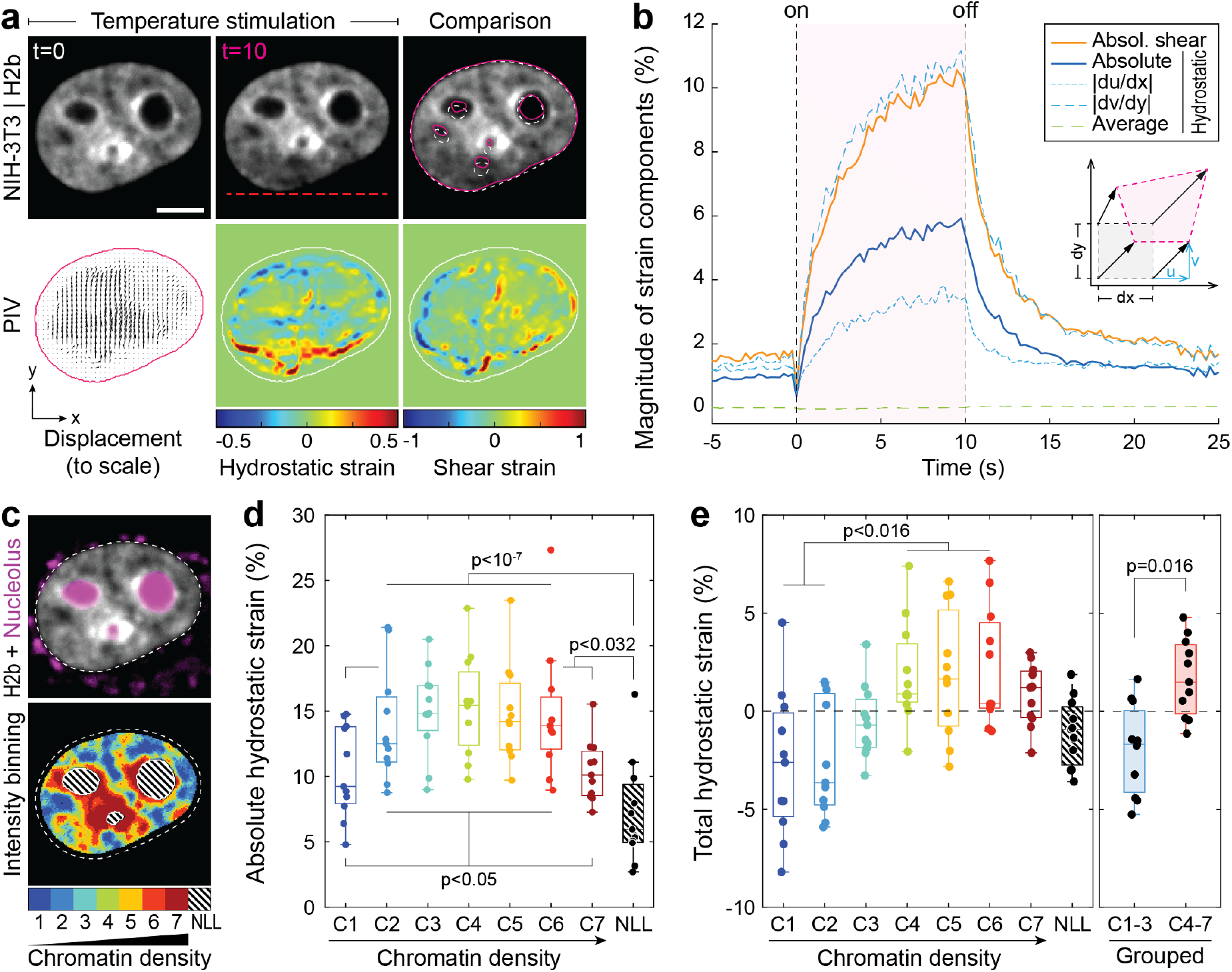
Differential strain measurement within chromatin compartments during temperature stimulation using spatially resolved strain maps. **a**) NIH-3T3 cells expressing mCherry-H2b were recorded during temperature stimulation. Spatially resolved displacement maps were generated from image stacks via PIV, using t=0s as undeformed reference frame. From this, hydrostatic and shear strain maps were further calculated. Strains are shown here as relative values (1=100%). Displacement map is shown at half density, see Suppl. Fig. 2 for full density and large-scale strain maps; scale=5 μm. **b)** Magnitude and temporal evolution of strain components. The inset cartoon depicts the concept of different strain types as a measure of local deformation. **c)** Nuclei were segmented into 7 equi-volumetric chromatin compartments of different density inferred by mCherry-H2b intensity. Nucleoli were further detected using Cytopainter live stains. A region of 6 pixels (∼ 0.74 μm) away from the nuclear border was cut off to exclude low displacements close to the nuclear lamina. **d)** Magnitudes of absolute hydrostatic strains were locally evaluated for different chromatin densities by combining strain maps with compartment maps. Shown is the averaged absolute hydrostatic strain for each compartment measured at peak deformation (t=10s) for n=11 nuclei. Boxplots depict the 25-75 percentile with whiskers spanning the full data range excluding outliers (>3xSTD). Statistics via one-way ANOVA with Tukey HSD. **e)** Local analysis of averaged total hydrostatic strains of n=11 nuclei showing that lightly packed chromatin bins (C1-3) are preferentially compressed while densely packed chromatin bins are extended. Statistics via one-way ANOVA with Tukey HSD for all groups and two tailed T-test for binned groups.

Using mCherry-H2b intensities as a proxy for chromatin density, we generated discretized maps of 7 nuclear compartments (Cremer et al., 2020) of equal volume (**Fig. 4c**). An additional dye was used to identify the nucleolus. Segmented fluorescence maps were then spatially correlated with strain maps to investigate whether there are differences in compartment behavior during temperature-induced chromatin displacements. Surprisingly, analysis of absolute hydrostatic strains revealed a non-linear relationship over chromatin densities, with most dense (C7) and most light packed regions (C1) experiencing the least, and medium dense regions (C4) the highest change in volume (**Fig. 4d-e, Suppl. Fig. 3**). Despite similar propensities in volume change, analysis of total (non-absolute) hydrostatic strains further showed that lightly packed euchromatin bins (C1-3) tend to be compressed, while heterochromatin bins (C4-7) tend to be extended (**Fig. 4f**). This likely reflected the inability of densely compacted chromatin to be further compacted, while loosely packed euchromatin seems to act as a mechanical buffer for decompaction inside a conserved volume.

The nucleolus is considered to be a liquid condensate that, despite lacking a membrane, maintains its integrity through liquid-liquid phase separation (Lafontaine et al., 2021; Strom and Brangwynne, 2019). Surprisingly, nucleoli showed high mechanical resilience during temperature induced chromatin displacement, as we measured only half the amount of absolute hydrostatic strain compared to overall chromatin (7.2 vs 13.5%) and a third less compared to dense chromatin regions (C7, 7.2 vs. 10.4%) that are frequently found adjacent to nucleoli.

### Immobile nucleoli provide a model case to study chromatin-nucleoli and chromatin-chromatin interactions

The nucleolus is formed by nucleolar organizing regions (NORs) that encode for different ribosomal RNAs (Bersaglieri and Santoro, 2019). These regions form a characteristic ring of condensed chromatin around the nucleolus, referred to as perinucleolar chromatin. In contrast to the lamina, the molecular mechanisms of chromatin-nucleoli tethering are not well understood (Mirny and Dekker, 2021). We observed that the nucleus showed high mechanical resilience to deformation. Moreover, in some cases, we observed that the nucleoli remained largely static during our stimulations with chromatin appearing to flow around it (**Fig. 5a, Suppl. Video 5**). To gain more insights into the way the nucleolus is mechanically embedded in nucleoplasm, we quantified the displacement of nucleoli and of adjacent 4 pixel thick (∼0.5 μm) perinuclear regions with a distance of 0-0.5 μm (PC1) 0.5-1.0 μm (PC2) and 1.0-1.5 μm (PC3) from the nucleoli border. Quantification verified that nucleoli (NLL), which appeared static during temperature induced chromatin displacement, moved significantly less compared to the nuclear average (NUC, **Fig. 5b**). While chromatin appears to flow around nucleoli in videos and PIV displacement maps, detailed analysis showed that the displacement of the closest regions (PC1) was not significantly higher than that of nucleoli (**Fig. 5c**). Displacements increased successively between the perinucleolar regions, but were still distinctly lower in PC3 compared to average chromatin displacements (NUC). This further suggested that, first, perinucleolar chromatin has a strong association with the liquid interface of the nucleoli border and, second, that the chromatin network is highly interwoven with signatures of continuous interaction on the scale of microns. As a control, velocity gradient does not occur around mobile nucleoli (**Suppl. Fig. 4**).

**Fig 5:**
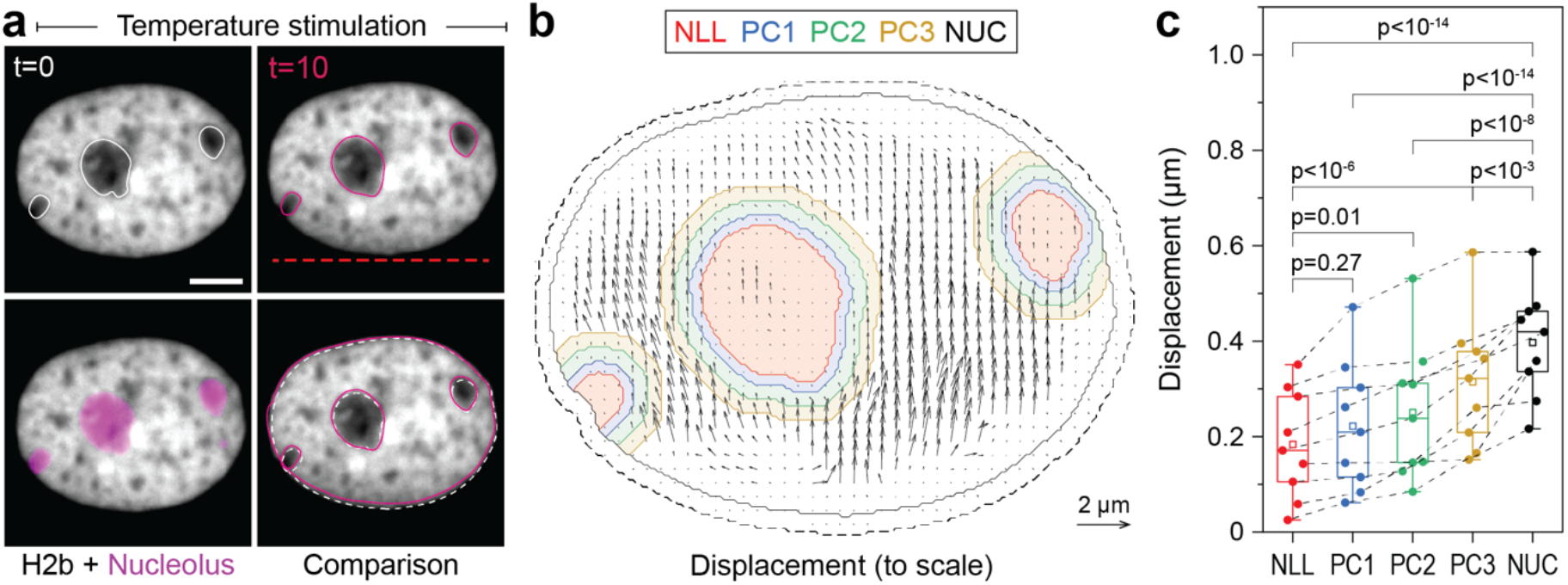
Measurement of chromatin displacements around static nucleoli reveal robust binding of perinuclear chromatin to the nucleolus as well as chromatin network effects. **a)** NIH-3T3 cells expressing H2b-GFP were stained with a nucleoli live stain and recorded during temperature induced chromatin displacement. Outlines represent detected nucleoli. Shown is an example in which nucleoli displayed little displacement, with chromatin flowing around the nucleoli like an obstacle. See also Suppl. Video 5; scale=5 μm. **b)** Displacement maps, derived via PIV, show chromatin motion around an immobile nucleolus. Indicated are the outlines of the nucleolus (NLL) and perinucleolar chromatin shells with a distance of 0-0.5 μm (PC1), 0.5-1 μm (PC2) and 1-1.5 μm (PC3) as well as the nuclear border (NUC). Low displacements close to the nuclear lamina (6 pixels ∼ 0.74 μm, dotted line) were excluded to better reflect internal chromatin motion. Displacement maps are shown at half density. **c)** Displacements for the nucleolus (NLL), perinuclear chromatin shells (PC1-3) and the nucleus as a whole (NUC) were quantified for n=9 different nuclei that displayed static nucleoli. See also Suppl. Fig. 4 for the cases of moving nucleoli as a comparison. Statistics via one-way ANOVA with Tukey HSD.

## DISCUSSION

We demonstrated that strongly localized temperature gradients can move chromatin non-invasively inside cell nuclei, thereby providing a complementary approach to existing methods to gain detailed new insight into the spatial organization of the nucleus. By displacing chromatin in a model of nuclear inversion (Mattar et al., 2018; Solovei et al., 2009), we observed high rigidity of the centrally formed heterochromatin cluster despite large displacements. This provided further evidence that highly compacted heterochromatin does not behave like a liquid but rather like a solid at micron scale, as recently suggested (Strickfaden et al., 2020). More general, in conventional interphase nuclei, we found that chromatin motion was largely reversible and showed exponential asymptotic trends over time. The simplest model that fit the deformation and relaxation dynamics was a Kelvin-Voigt model that features a viscous dash-pot and an elastic spring element connected in parallel, indicating that the underlying mechanisms of liquid and solid behavior are interwoven. The characteristic times τ extracted from this model where around 1 s. Since τ reflects the ratio of viscosity to elasticity (τ = η/E), this suggested that liquid and solid contributions of chromatin, specifically the phases C1-C6, are in close balance on the mesoscale when assessed on the time scale of seconds.

We further showed that different nuclear compartments possess distinct material properties. Specifically, we found that the susceptibility to volumetric deformation (absolute hydrostatic strain) showed a non-linear relationship over chromatin compaction with medium dense chromatin being most compliant, and lightly and densely packed chromatin being most resistant to deformation. A similar non-linear relationship between hydrostatic strain and chromatin density has been reported in cardiomyocyte nuclei during spontaneous cell contractions (Ghosh et al., 2019). That lightly packed chromatin compartments show high resilience is still somewhat surprising and merits further investigation. One reason could be that persistent tethering of RNA to transcribed chromatin, important for the formation of transcriptional pockets, provides structural support (Hilbert et al., 2021).

Furthermore, we observed a striking mechanical identity of the nucleolus, which showed higher resistance to deformation than any chromatin compartment. The nucleolus is considered to be a membraneless liquid droplet. Our findings might help to better understand the underlying physics (e.g. its surface tension) that allow the nucleolus to maintain a stable form (Caragine et al., 2019; Feric et al., 2016). We frequently observed that the nucleolus resisted displacement altogether. A reason for that could be that perinucleolar chromatin further anchors nucleoli to the nuclear lamina, especially in 2D cultures where nuclei have a flat topology (Ghosh et al., 2019). In such cases we found that perinucleolar chromatin showed similar resistance to displacement, suggesting a tight association to the nucleolus. Nucleolus associated domains (NADs) are thought to anchor peri-nucleolar heterochromatin to the nucleolus (Canat et al., 2020), but more needs to be understood.

Based on recent FRAP experiments that revealed that chromatin does not mix and recover, while chromatin scaffold proteins rapidly do, the authors suggested that interphase chromatin is akin to a porous hydrogel. Our results support this view, and complement the FRAP based evidence by detailed analysis of the mechanical relaxation response after perturbation. Specifically, our method reveals a characteristic dynamic behavior of a porous gel-like phase with both dissipative as well as elastic contributions, the latter of which being responsible for deformations being predominantly reversible. When analyzing chromatin motion around static nucleoli, one can directly observe that chromatin shows ‘network effects’, meaning coherent motion of spatially extended gel-like heterochromatin domains. These network effects become visible over length scales of up to 1.5 μm as a smooth gradient of velocities surrounding immobile nucleoli and are consistent with coherent motion and relaxation of genetic loci after displacement by magnetic forces (Keizer et al., 2021).

As different methods shed light onto different aspects of nuclear organization, combining our approach with other complementary methodologies will be useful to reach an integrated view of nuclear organization. For example, combining data from live perturbations with chromatin conformation capture methods (Hi-C) might be key to connect mechanical identities of compartments with their underlying sequence interactions (Hildebrand and Dekker, 2020). Specifically, our method could be used in conjunction with the recently developed liquid Hi-C approach that aims to disentangle the contribution of the chromatin backbone and non-covalent chromatin interactions for nuclear mechanics and organization (Belaghzal et al., 2021). Similarly, ChIP-seq approaches could be employed to further elucidate the roles of epigenetic modifications and chromatin-protein binding in shaping these interactions (Huang et al., 2015; Jiang and Mortazavi, 2018; Mourad and Cuvier, 2015).

We also showed that this method allows to study the material interfaces between compartments, such as chromatin and the nucleoli. Similar approaches could be used to study the interaction of chromatin with the nuclear lamina, especially to study diseases in which lamina dysfunctions cause aberrant nuclear organization, referred to as Laminopathies (Isermann and Lammerding, 2014; Köhler et al., 2020; Stiekema et al., 2020). Of high interest would also be to study the transition of this lamina-interaction during mitosis to achieve a better understanding of the underlying mechanism of nuclear reformation (Serra-Marques et al., 2020).

While our method constitutes a reliable, non-invasive and well tunable way to induce chromatin motion in cell nuclei and study its relaxation behaviors, our perturbations bear further potential to gain insight into the physical chemistry that underlies temperature dependent chromatin organization. Temperature dependent changes in chromatin compaction have been reported after bulk cooling of live cell nuclei (Fischl et al., 2020), albeit on the time scale of hours. On shorter time scales, reversible changes in nuclear volume have been observed after homogenous temperature increases (ΔT=18°C) in isolated nuclei (Chan et al., 2017). Interestingly, the study found that the sign of volumetric change was dependent on ion valency, especially multivalent cations, hinting towards an electro-osmotically driven influx of water into these isolated nuclei. As the directed motion of chromatin within a cell nucleus as described by us occurs without signs of such nuclear volume changes and at about ten-fold smaller temperature differences (2 vs. 18°C), it is likely driven by the temperature gradient.

A wide range of physical phenomena is known that give rise to the motion of microscopic objects in temperature gradients. The movement of molecules along a temperature gradient (thermophoresis or Soret effect) is complex and subject of ongoing scientific debates. However, studies have shown that the movement of highly charged polymers, such as DNA and RNA, in aqueous solutions can be predicted over a large range of experimental parameters by the temperature gradient-induced emergence of local and global electric fields that link ionic thermophoresis to electrostatic energies and the Seebeck effect respectively (Duhr and Braun, 2006; Reichl et al., 2014).

Moreover, the observed actuation of chromatin could in parts also be driven by a temperature dependent affinity of DNA to histone complexes, potentially leading to a decompaction of condensed chromatin with increasing temperature. Equally, a temperature dependent hydrophilicity of chromatin, as it is known for certain polymers (Quesada-Pérez et al., 2011) and has technologically been exploited to elicit responses phenomenologically similar to the bending of bi-metallic strips (Hippler et al., 2019), could potentially give rise to chromatin motion in temperature gradients. Additionally, motion in fluids may also be the result of so-called thermoviscous flows (Weinert et al., 2008). These have successfully been used to stream the cytoplasm (Mittasch et al., 2018), but require the spatial scanning of a temperature field, and as such can be decoupled from the here observed effects due to their fundamentally different symmetry properties of the stimulus.

In conclusion, we showed that strongly localized temperature gradients offer unexpected opportunities to study the organization of the living nucleus in a spatially resolved and dynamic manner.

## MATERIALS AND METHODS

### Cell Culture and Transfection

NIH-3T3 cells were cultured in DMEM + GlutaMAX (Gibco) containing 10% fetal bovine serum (Gibco) and 1% penicillin-streptomycin (Gibco) at 37°C and 5% CO2. For temperature stimulation experiments, 150 μm thick c-axis cut sapphire cover slips (UQG Optics) were coated with fibronectin (60 μg/mm^2^) for 1h at RT and seeded with cells to reach 50% confluency the next day. Sapphire was chosen for its excellent heat conductivity while still allowing for high quality imaging. Transfection of GFP-H2b, mCherry-H2b or Casz1_v2 (NM_017766) containing plasmids was performed 18h after seeding using Lipofectamine 3000 and cells were incubated for another 24h before experiments. To visualize nucleoli, cells were stained with Cytopainter Nucleolar Staining Kit (Abcam, ab139475) 30 min before experiments. During experiments, a temperature chamber, consisting of a thick sapphire glass slide with Peltier elements on each side (Mittasch et al., 2018), was used to maintain a constant ambient temperature of 36°C (Peltier elements convert heat into energy and vice versa). On the day of experiments, sapphire cover slips containing transfected cells were mounted onto temperature chambers using 15 μm polystyrene spacer beads (Bangs Laboratories).

### Live Cell Imaging and Temperature Stimulation

Image stacks were taken on an inverted Olympus IX81 microscope equipped with a Yokogawa spinning disk confocal head (CSU-X1), 60x 1.2 NA plan apochromat water objective and an iXon EM + DU-897 BV back illuminated EMCCD (Andor). Images were acquired at 4 frames per second, with an excitation of about 200ms, using VisiView software (Visitron Systems). Cells were imaged for 1 min total, starting with 20s of no stimulation (baseline), followed by 10s of temperature stimulation and ending with 30s of no stimulation again (reversibility).

To apply a local, precisely controlled temperature gradient, an infrared laser (1455 nm) was scanned along a line at 1 kHz. The exact setup has been described before (Mittasch et al., 2018). Briefly, an infrared Raman laser beam (CRFL-20-1455-OM1, 20 W, near TEM00 mode profile, Keopsys) was acousto-optically scanned along a line. Precise deflection patterns were generated using a dual-channel frequency generator PCI card (DVE 120, IntraAction), controlled via modified LabVIEW (National Instruments) based control software (DVE 120 control, IntraAction), in combination with a power amplifier (DPA-504D, IntraAction). For two-dimensional laser scans (Fig. 4a), a two-axis acousto-optical deflector (AA.DTSXY-A6-145, Pegasus Optik) was used. Precise laser scan patterns were performed by generating analog signals using self-written software in LabVIEW, in combination with a PCI express card (PCIe 6369, National Instruments). A dichroic mirror (F73-705, AHF, Germany) was used to couple the infrared laser beam into the light path of the microscope by selectively reflecting the infrared light but transmitting visible wavelengths which were used for fluorescence imaging.

### Dye-Based Temperature Measurements

To measure the spatial temperature profile inside the cell incubation chamber, as well as the temperature increase inside nuclei during temperature stimulation experiments, we used temperature sensitive decrease in quantum efficiency that has been well-described for certain dyes (usually in the red spectrum) before (Hirsch et al., 2018; Mittasch et al., 2018; Singhal and Shaham, 2017). For chamber measurements, Rhodamine B solution (Sigma, 02558) was diluted to 10% in water and image stacks were acquired in the red channel during laser application. For nuclear measurements, mCherry-H2b transfected cells were recorded. Both dyes were calibrated by precise changing of the bulk temperature of the incubation chamber (Suppl. Fig. 1). For Rhodamine, thermophoretic effects (lower dye intensity due to concentration difference, not quantum yield) were determined to correct measurements.

### Displacement and Strain Map Calculation

A custom MATLAB code was written to calculate spatial displacements and strain maps from image stacks. A modified version of MatPIV (v 1.7) (Sveen, 2004) was used to generate displacement maps with a window size of 32 pixels, 75% overlap using multiple passes as well as local and global filters. The final displacement field resolution was 4×4 pixels and displacement maps in this manuscript are shown only at half resolution. From displacement maps, hydrostatic and shear strain maps were calculated according to:

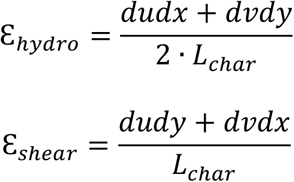

With L_char_ being the characteristic (initial) length before deformation.

To extract local strain information, binary masks of chromatin densities, using intensity histograms, or of nucleoli regions using intensity thresholding were generated. Strain maps were extrapolated to match image resolution and local strains were averaged using binary masks. The same algorithm was used to track changes in nuclear and CHC area.

Image difference stacks and average image difference tracks were generated using ImageJ (v. 1.52t). The “Analyze Particles” function in ImageJ was used to track the centroid position of nuclear features and calculate feature displacements. Dynamic measurements of image differences and displacements were fitted to a Kelvin Voigt Model, consisting of a spring and a dashpot in series, using the equations:

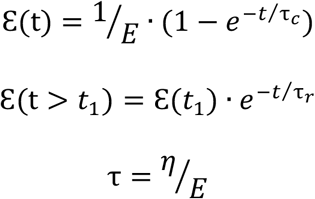

For the creep (during stimulation) and relaxation phase (after stimulation), respectively, with E being the relative spring constant and τ the characteristic time, which reflects the ratio of viscosity (η) to elasticity (E) (Meyers and Chawla, 2009).

### Statistics

T-tests or One-Way ANOVA were conducted using OriginPro 2021 (v. 9.8.0.200). The statistical test used, as well as the number of repeats n and significance levels, are indicated in the figures and/or in the figure captions. All data with repeated measurements were collected in at least 3 independent experiments.

## ACKNOWLEDGEMENTS

We want to thank Falk Elsner and Claudius George for constructing the temperature incubation chambers. Further, we would like to thank Irina Solovei, Job Dekker and Denis Lafontaine for discussions as well as Lennart Hilbert and Iain Patten for feedback on the manuscript, and Anatol Fritsch and Matthäus Mittasch for devising the setup used for the thermal manipulations.

## AUTHOR CONTRIBUTIONS

Conceptualization, B.S., and M.K.; Methodology, B.S., S.K. and M.K.; Software, B.S. and S.K.; Formal Analysis, B.S. and M.J.; Investigation, B.S., M.J, E.E. and I.S.; Interpretation of results, B.S., I.S. and M.K, Writing – Original Draft, B.S., I.S. and M.K.; Writing – Review & Editing, All authors; Funding Acquisition, M.K.;

## SUPPLEMENTAL MATERIAL

**Suppl. Fig 1:**
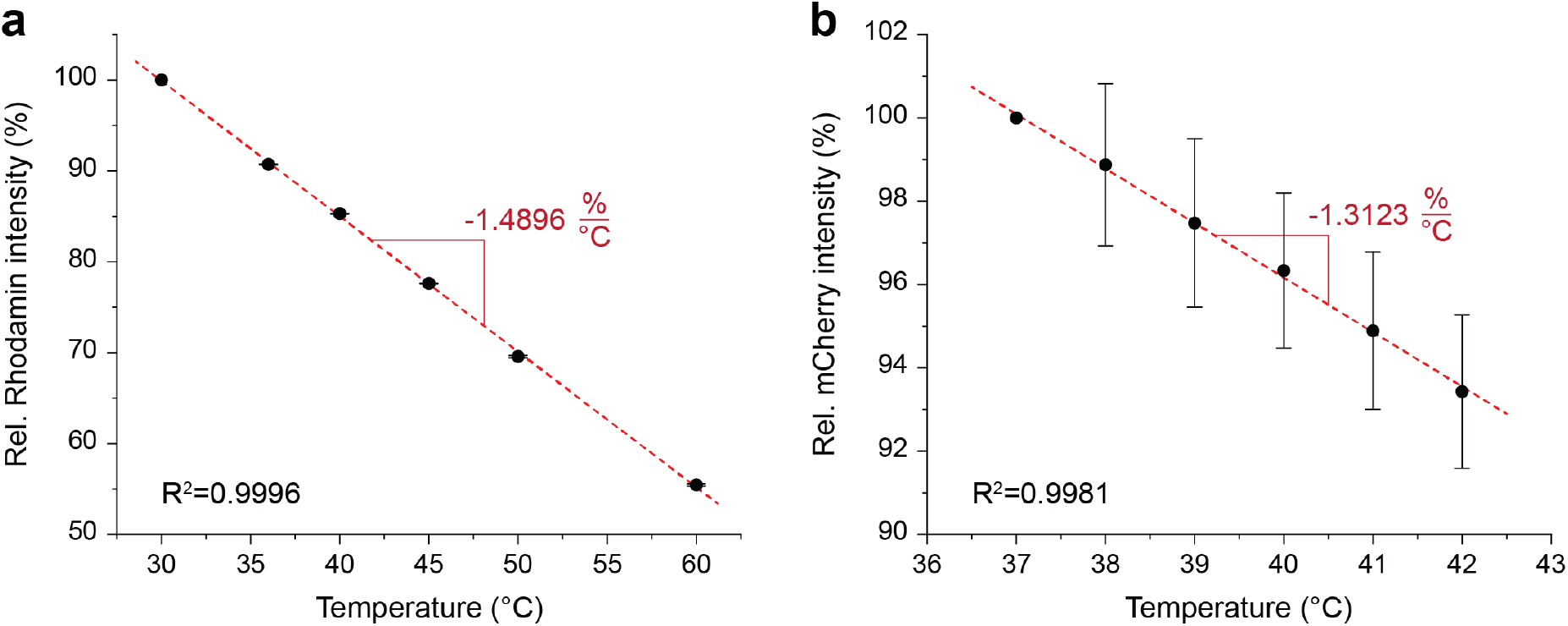
Calibration of dyes for temperature measurements. Quantum efficiency (relative intensity) of **a)** Rhodamine and **b)** mCherry was measured at different temperatures via the same temperature incubation chamber used for live cell experiments; n=5, error=STD.

**Suppl. Fig 2:**
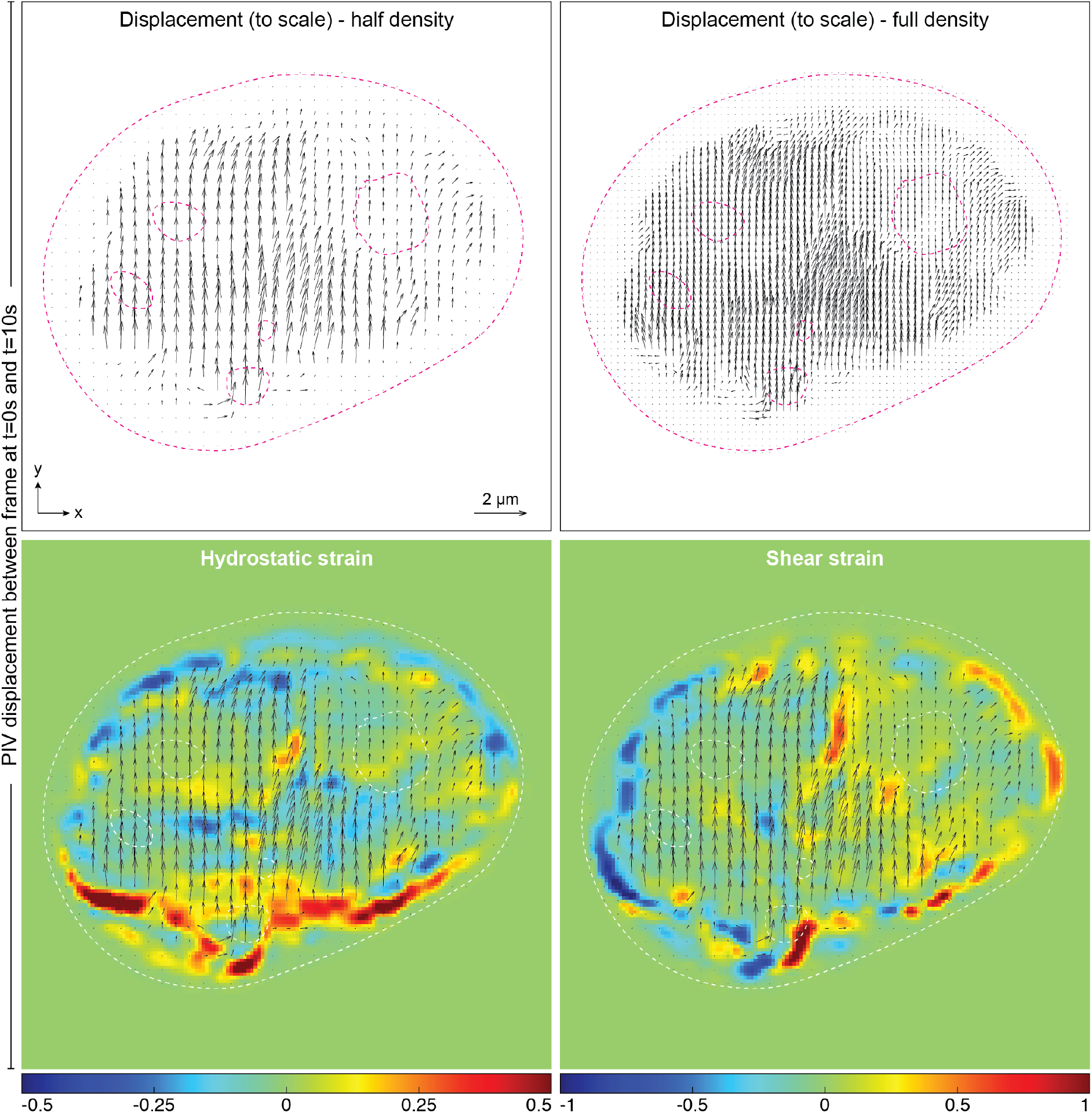
Displacement and strain maps. Large scale images of displacement maps, generated via PIV, and derived strain maps between rest (t=0s) and peak displacement (t=10s) corresponding to Fig. 4a.

**Suppl. Fig 3:**
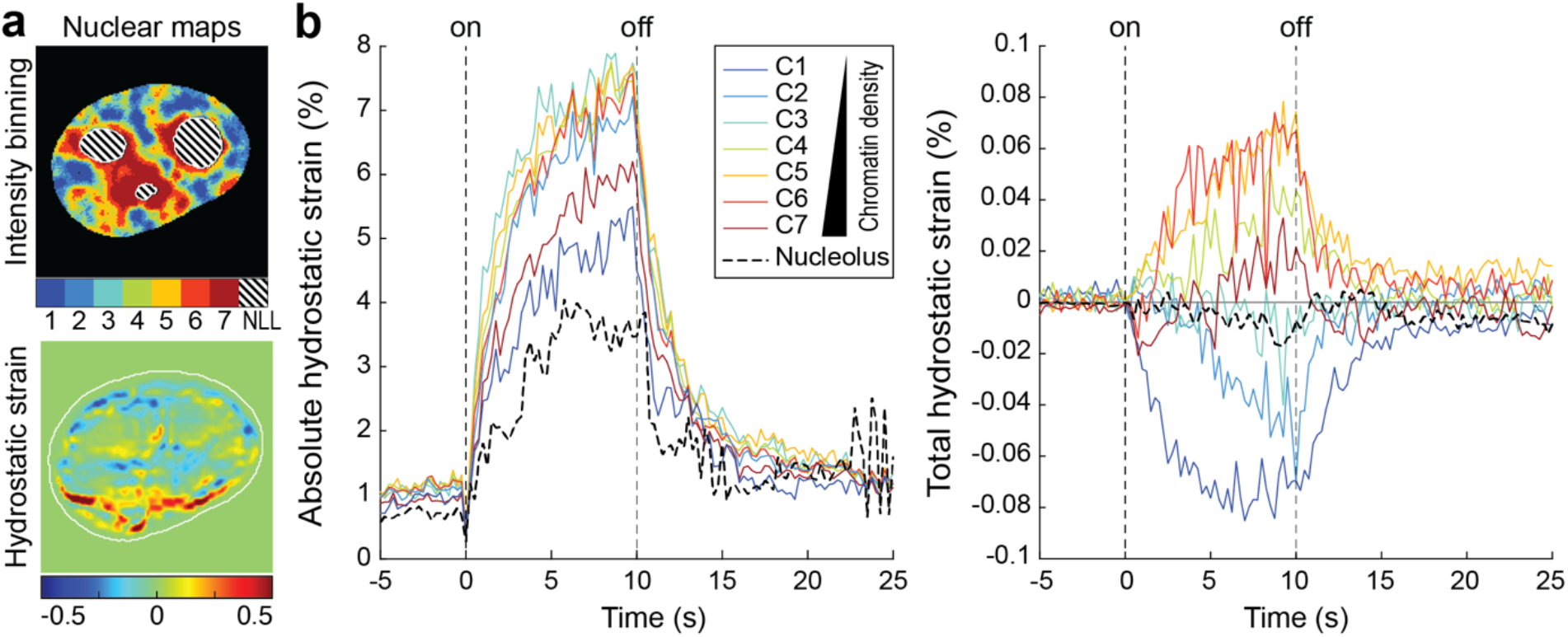
Hydrostatic strain over time for different nuclear compartments of a single nucleus. **a)** Top: Nuclear compartments with different chromatin densities as well as nucleoli were mapped via mCherry-H2b and a nuclear live stain. Bottom: Intra-nuclear strain maps were calculated from chromatin displacements, derived via PIV. **b)** Compartment and strain maps were combined to analyze mechanical identities of different chromatin compartments. Shown are absolute and total hydrostatic strains of different nuclear compartments (C1-7) and the nucleolus (NLL) over time.

**Suppl. Fig 4:**
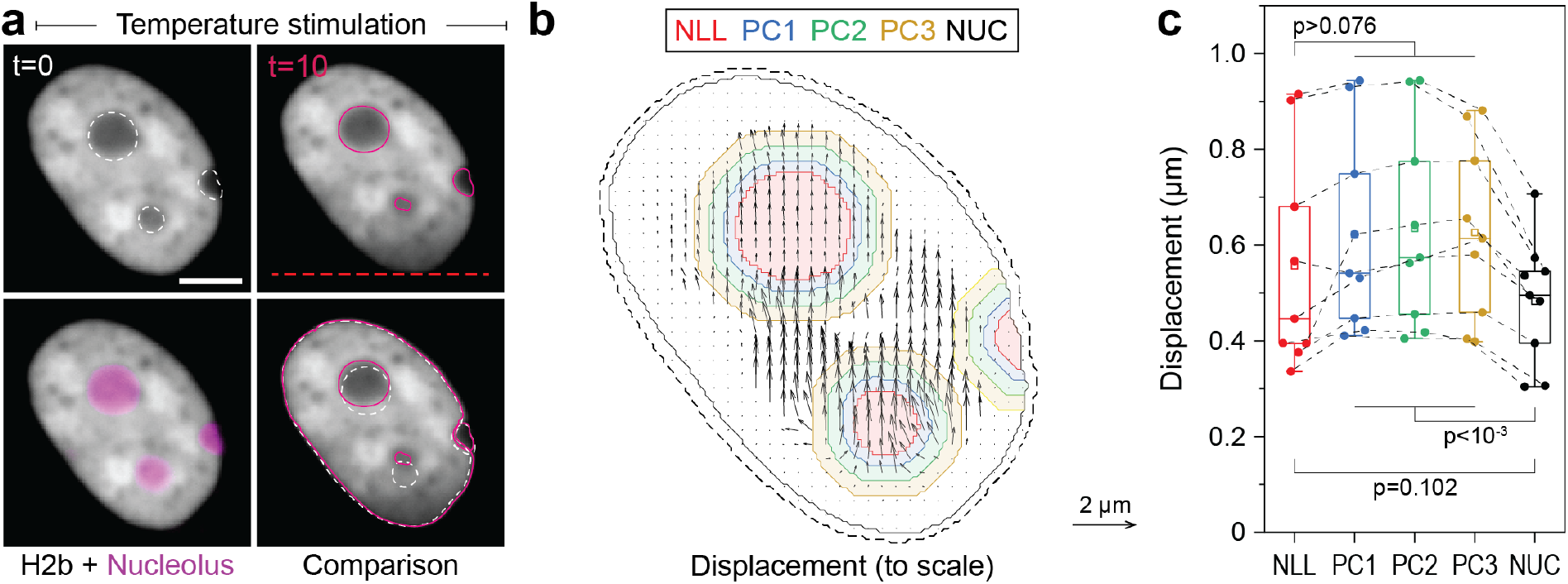
Displacement analysis of moving nucleoli as comparison for static nucleoli. **b)** NIH-3T3 cells expressing H2b-GFP were stained with a nucleoli live stain and recorded during temperature induced chromatin displacement. Outlines represent detected nucleoli. Shown is an example in which nucleoli showed displacements similar to surrounding chromatin; scale=5 μm. **b)** Displacement maps were derived via PIV. Indicated are the outlines of the nucleolus (NLL) and perinucleolar chromatin shells with a distance of 0-0.5 μm (PC1), 0.5-1 μm (PC2) and 1-1.5 μm (PC3) as well as the nuclear border (NUC). Low displacements close to the nuclear lamina (6 pixels ∼ 0.74 μm, dotted line) were excluded to better reflect internal chromatin motion. Displacement map is shown at half density. **c)** Displacements for the nucleolus (NLL), perinuclear chromatin shells (PC1-3) and the nucleus as a whole (NUC) were quantified for n=9 different nuclei that displayed moving nucleoli. Statistics via one-way ANOVA with Tukey HSD.

**Suppl. Video 1-2: Video microscopy of temperature stimulated and control nuclei, corresponding to image difference analysis in Fig. 1**. GFP-H2b transfected NIH-3T3 cells were recorded during stimulation with a temperature gradient, placed along the bottom of the nucleus (Video 1), or without stimulation (Video 2). Recorded at 4 fps, show at 4x real-time (16 fps). Stimulation for 10s.

**Suppl. Video 3: Video microscopy of temperature stimulated nuclei, corresponding to feature tracking analysis in Fig. 2**. A GFP-H2b transfected NIH-3T3 cell was recorded during stimulation with a temperature gradient placed along the bottom of the nucleus. Recorded at 4 fps, show at 4x real-time (16 fps). Stimulation for 10s.

**Suppl. Video 4: Video microscopy of temperature stimulated inverted nuclei, corresponding to analysis in Fig. 3**. A Casz1 and GFP-H2b double-transfected NIH-3T3 cell that formed a centrally compacted heterochromatin feature was stimulated with a temperature gradient placed along the bottom of the nucleus. Recorded at 8 fps, show at 2x real-time (16 fps). Stimulation for 5s.

**Suppl. Video 5: Video microscopy of temperature stimulated nuclei, showing a case of static nucleoli and corresponding to analysis in Fig. 5**. A GFP-H2b transfected NIH-3T3 was stained with a nucleoli specific dye and was recorded during stimulation with a temperature gradient placed along the bottom of the nucleus. H2b (white) and nucleoli (magenta) channels were recorded sequentially at 2 fps, shown at 8x real-time (16 fps). Stimulation for 10s.

